# Homology-Based Variant-Effect Predictors Break Down on Cytochrome P450 Pharmacogenes^*^

**DOI:** 10.64898/2026.07.30.741616

**Authors:** Helen Xu, Issah Samori, Gowri Nayar, Russ B. Altman

**Affiliations:** School of Medicine, Stanford University, Stanford, CA, USA; Department of Bioengineering, Stanford University, Stanford, CA, USA; Department of Biomedical Data Science, Stanford University, Stanford, CA, USA; Department of Genetics, Stanford University, Stanford, CA, USA

**Keywords:** Cytochrome P450, variant-effect prediction, pharmacogenomics, protein language models, deep mutational scanning

## Abstract

Cytochrome P450 (CYP) enzymes metabolize roughly three-quarters of clinically used drugs, and genetic variation in these enzymes is a leading source of interindividual differences in drug response. Predicting a variant’s functional effect is therefore critical, yet the consequences of most CYP variants remain unknown. Many state-of-the-art variant-effect predictors rest on a homology-based paradigm that scores variants by evolutionary conservation — an assumption that pharmacogenes, including CYPs, violate. Focusing on human CYPs, we show that homology-based models fail systematically. AlphaMissense (AM) assigns variants to its “ambiguous” class at nearly twice the proteome-wide rate across six clinically important CYPs, and within that class its scores are essentially uninformative about measured CYP2C9 deep mutational scan (DMS) activity (*ρ* = −0.130). Using catalytic activity and cellular abundance from the same CYP2C9 DMS, we show that AM is largely an abundance signal: it tracks activity well for destabilizing, low-abundance variants (*ρ* = −0.568) but far more weakly for abundance-independent variants (*ρ* = −0.357; Fisher *r-to-z z* = −8.99, *p* < 10^−18^). We further hypothesized that non-homology-based features (sequence position, substitution chemistry, distance to the substrate-recognition sites, and secondary structure) might account for where AM errs, but their explanatory power is weak. Using CYP2C9 DMS activity as ground truth, we built a *k*-nearest-neighbors model over ESM-2 embeddings and ensembled it with AM and ESM-2 masked marginal probability, raising the correlation between predicted and measured activity within the ambiguous class from *ρ* = 0.080 for AM alone to 0.399, and reaching an overall *ρ* = 0.721 that exceeds every out-of-the-box predictor we tested (*ρ* = 0.523 to 0.686). However, neither this improved ensemble model nor any other method agrees significantly with clinical annotations. Drawing on evidence that a variant’s effect can depend on the drug, we hypothesize that substrate identity is the key missing feature in current models, and that predicting function for these multi-substrate enzymes may require redefining function as substrate-conditioned.

## 1. Introduction

Adverse drug reactions are among the leading causes of death in hospitalized patients. Much of this harm is not idiosyncratic but genetic: interindividual differences in the enzymes that metabolize drugs mean that the same dose can be therapeutic for one patient and fatal for another.^1,2^ Chief among these enzymes are the cytochrome P450s (CYPs), which are involved in the metabolism of roughly three-quarters of all clinically used drugs.^3–5^ Genetic variation in CYP genes is correspondingly one of the largest sources of interindividual variability in drug response.^6,7^ CYP2C9 alone metabolizes up to *∼*15% of small-molecule drugs, including the widely prescribed anticoagulant warfarin, for which variant-driven differences in metabolism directly affect safety and efficacy.^8^ Pharmacogenomic testing and genotype-guided prescribing aim to pre-empt these outcomes,^9,10^ but in practice they rest on a small set of well-characterized “star” alleles. The functional consequences of the vast majority of observed CYP variants remain unknown, so we generally cannot predict whether a given variant will increase, decrease, or leave unchanged the metabolism of a specific drug. Closing this prediction gap is a prerequisite for scaling pharmacogenomics beyond its current handful of actionable alleles. The number of possible variants vastly exceeds what deep-mutational, biochemical, or clinical characterization can hope to interrogate, so filling this gap necessarily requires computational variant-effect prediction. Yet most widely used predictors annotate a variant by its similarity to sequences shaped by evolution — the same homology-based paradigm that computational function annotation has relied on for decades. SIFT aligns related sequences and scores a substitution by how often the mutant residue is tolerated at that position;^11^ AlphaMis-sense (AM) couples AlphaFold with multiple-sequence-alignment (MSA) input and human and primate allele frequencies to predict pathogenicity against ClinVar benchmarks;^12,13^ protein language models of the ESM family absorb the same signal by masked-language modeling over divergent sequences, whether repurposed zero-shot through masked marginal probabilities (MMP), as in ESM-2,^14,15^ or purpose-built for variant-effect prediction, as in ESM-1v.^16^ Three further model families build on this idea: REVEL is a supervised ensemble that combines conservation- and structure-derived component scores against pathogenicity labels;^17^ DeMaSk is fitted to a substitution matrix derived from deep mutational scans and is therefore trained on measured function rather than on conservation alone;^18^ and PharmGScore is tuned specifically for pharmacogene variants, the setting where generic predictors are known to degrade.^19^

We use *homology-based* throughout for predictors whose variant-effect signal derives, explicitly or implicitly, from the distribution of residues observed at a position across related sequences — retrieved at inference by alignment, as in SIFT and AM; absorbed during pre-training, as in ESM-2; aggregated from component scores that are themselves alignment-derived, as in REVEL; or supplied as population allele frequencies, as in other related models. This definition is deliberately broad, and it includes the protein-specific model we build here. The premise that evolutionary tolerance is a good proxy for functional consequence works well for the disease genes these models are usually tuned on: such genes are under strong evolutionary pressure to stay unchanged, so a variant that breaks conservation is likely to be harmful. Yet pharmacogenes violate that assumption. CYPs evolved largely to metabolize compounds in the diet and environment, and their central role in drug metabolism is in a sense incidental to that history.^20,21^ As a result, reduced or even absent activity of an individual CYP is frequently benign on its own, with consequences that surface only when a particular food or drug is encountered. CYP variants are therefore usually not disease-causing; they frequently occur at high population frequencies; and they arise in a gene family that is unusually polymorphic and functionally diverse rather than strongly conserved.^4,6^ Consistent with this mismatch, homology- and pathogenicity-oriented predictors perform markedly worse on pharmacogenetic than on disease variants, motivating a growing body of pharmacogene-specific methods and evaluations.^19,22–25^ What has remained unclear is whether this degradation reflects mere low model confidence or a genuine information ceiling, in which an important determinant of function is missing and cannot be recovered by better calibration alone.

Among pharmacogenes, CYPs are one of the more tractable settings in which to examine this question, combining the violated assumptions with clinical necessity and a rich, though still incomplete, body of ground truth. Decades of research have produced variant-level data spanning high-throughput deep mutational scanning (DMS) of enzymatic activity and abundance,^8,26^ clinical annotations, and curated pharmacogenomic databases, yet the functional consequences of most CYP variants are still unresolved. Because CYPs are comparatively well characterized, cases where homology-based predictors fail more plausibly expose a limit of the homology paradigm than an absence of data.

In this work, we use human CYP pharmacogenes as a case study for where and why homology-based variant-effect prediction breaks down. Our contributions are:

1. We show that AM assigns CYP variants to the “ambiguous” class at nearly twice the proteome-wide rate across six CYPs, and that on CYP2C9 — the one CYP with paired activity and abundance DMS — its activity signal is abundance-based and weak on the abundance-independent, catalytic axis, particularly within the ambiguous class, with structural and biochemical priors accounting for almost none of the residual error.
2. We show that a *k*-nearest-neighbors model over ESM-2 embeddings, ensembled with AM and ESM-2 MMP, outperforms generic, ensemble, function-trained, and pharmacogene-tuned predictors on CYP2C9 activity, yet this accuracy does not translate into reliable agreement with clinical or pharmacogenetic labels.
3. We propose substrate identity as the leading candidate for the missing, non-homologous variable, drawing on pharmacogenetic evidence that a single variant’s consequence can depend on the drug, and argue that substrate-conditioned prediction is a promising route beyond homology.

Taken together, these results reframe the annotation problem for pharmacogenes: the barrier is not simply that homology-based similarity is too weak, but that the definition of function these models predict — a single substrate-agnostic effect — is the wrong target for multi-substrate enzymes. Moving beyond homology here may therefore require redefining function itself, as a quantity conditioned on substrate, rather than more or better similarity search.

## 2. Methods

### 2.1. Datasets and variants

We drew on four ground-truth resources, each encoding a distinct notion of variant effect. DMS activity measurements were taken from Amorosi et al. (2021),^8^ which reports click-seq (enzymatic activity, *N* = 6,273) and vamp-seq (cellular abundance, *N* = 6,559) scores for CYP2C9 missense variants, with *n* = 4,478 overlapping variants. We used the normalized click-seq activity score as the primary quantitative target throughout and the abundance readout for the abundance-stratification analysis (Section 2.3.2). ClinVar provided disease-oriented clinical significance labels following the ACMG framework;^27,28^ after filtering for CYP2C9 single-missense variants with a functional classification, 50 variants at 42 unique positions remained. PharmVar provided pharmacogenetic function classes;^29^ we used the 77 CYP2C9 star alleles defined by a single missense substitution with functional classification. ClinPGx (PharmGKB) provided literature-curated variant–drug annotations for the substrate-resolved analysis (Section 2.3.5).^30^

The enrichment analysis (Section 2.3.1) and the substrate-resolved analysis (Section 2.3.5) spanned six clinically important CYPs (CYP1A2, CYP2C19, CYP2C9, CYP2D6, CYP2E1, and CYP3A4) whereas activity-benchmarked modeling focused on CYP2C9, for which comprehensive DMS data were available. Reference protein sequences for all CYPs were obtained from UniProt.^31^ All analyses were done in Python 3.12.11 on the Sherlock computing cluster operated by Stanford Research Computing.

### 2.2. Variant-effect predictors and models

#### 2.2.1. Predictors and model selection

We adapted three predictors (AM, ESM-2, and KNN) for CYP2C9 activity prediction. For CYP2C9 (UniProt P11712) and the five additional CYPs, we obtained AM pathogenicity scores together with AM’s discrete class labels (likely benign, ambiguous, or likely pathogenic), which AM assigns by thresholding its predicted pathogenicity score. ESM-2 (`esm2_t33_650M_UR50D`) was used as a zero-shot variant-effect signal via its MMP (MMP = log *P* (*x*_*i*_ = mut | *x*_*\i*_)−log *P* (*x*_*i*_ = wt | *x*_*\i*_)), and as protein embeddings for the KNN model (Section 2.2.2). We selected ESM-2 over ESM-1v for the main analyses because ESM-2’s zero-shot scores tracked CYP2C9 DMS activity more closely (Fig. S1).

Because AM and ESM-2 MMP scores are not on the activity scale, we calibrated each to click-seq activity using monotonic regression. We compared isotonic and logistic calibrators by 5-fold out-of-fold (OOF) *R*^2^ and selected the better-performing option. For AM this was isotonic regression; the monotonicity constraint encodes the expected biological direction, in which higher pathogenicity corresponds to lower activity. For ESM-2 MMP it was a four-parameter monotone logistic function, constrained to be monotone increasing so that higher MMP corresponds to higher activity. Calibrators were fit within each training fold only.

We evaluated four external baselines, scored through the same evaluation pipeline: DeMaSk,^18^ PharmGScore,^19^ SIFT,^11^ and REVEL.^17^ PharmGScore values were taken from a precomputed release and mapped to the Ensembl canonical CYP2C9 transcript (ENST00000260682) by CDS coordinate; SIFT and REVEL scores came from dbNSFP v4.8.^32^ To avoid data leakage, no predictor was evaluated on data to which it was fit. SIFT is unsupervised, and DeMaSk, though trained on DMS data, predates the CYP2C9 scan; both were therefore included in the activity and the clinical-alignment analyses. REVEL and PharmGScore were trained on pathogenicity annotations that include ClinVar labels, so both are reported for activity prediction but excluded from the clinical-alignment comparison.

#### 2.2.2. CYP-specific KNN over ESM-2 embeddings

We built a KNN regressor predicting click-seq activity from ESM-2 embeddings. Each mutated CYP2C9 protein, also referred to as a variant, was represented by the final-layer, per-residue ESM-2 representation, pooled over all sequence positions (excluding the BOS/EOS tokens) to a 1,280-dimensional vector.

All cross-validation in this work was grouped by residue position (GroupKFold) to prevent leakage from near-duplicate observations on both sides of a split, so that no variant is ever predicted by a model that has seen another substitution at its own residue. The scheme is nested: an outer 5-fold position-grouped split for evaluation; within each outer training partition, an inner 4-fold position-grouped split that generates the base-learner meta-features for the ensemble; and, for KNN hyperparameter selection, a 3-fold position-grouped split within each outer training partition that scores every configuration in the grid. The hyperparameter configuration selected most often across outer folds (mean pooling, Euclidean distance, distance-weighted averaging, *K* = 30) was used in all subsequent analyses (Fig. S2). For the clinical comparisons (Section 2.3.4), ClinVar and PharmVar variants were excluded from model training at the position level.

#### 2.2.3. Ensemble model

We combined the calibrated AM activity, calibrated ESM-2 MMP activity, and KNN-predicted activity using a ridge-regression meta-learner on standardized features (`StandardScaler` followed by `RidgeCV` over *α ∈* [10^−3^, 10^3^]). All models were evaluated by outer 5-fold cross-validation, and we report OOF Spearman’s *ρ* against click-seq activity.

### 2.3. Statistical analyses

#### 2.3.1. Ambiguous-class enrichment across CYPs

For each of the six CYPs we retrieved all possible missense variants and their AM classes, and compared the fraction assigned to the ambiguous class against the proteome-wide baseline reported by the original AM paper.^12^ Significance was assessed with a one-sided binomial test.

#### 2.3.2. Abundance stratification

We split variants at the median abundance into a low-abundance (destabilizing) and a normal/high-abundance (abundance-independent) subset, and compared the model’s score– activity Spearman correlation between strata, both across the full dataset and within each AM class. Because abundance is itself correlated with predictor score, a difference between strata could in principle reflect a narrower score range in one of them; we therefore verified that score range, interquartile range (IQR), and standard deviation were matched between the low- and high-abundance strata. Differences between strata were tested with an independent-samples Fisher *r*-to-*z* test.

#### 2.3.3. Structural and biochemical priors

We asked whether four interpretable features explain AM’s prediction error, defined as the absolute residual between its calibrated activity prediction and the measured DMS activity. **(1)** *Sequence position* of the point mutation, correlated with error by two-sided Spearman. **(2)** *Substitution chemistry*: error compared across wild-type residue groups (nonpolar GAVLIMPFW, polar STCYNQ, positively charged RKH, negatively charged DE) by Kruskal–Wallis test with *η*^2^ as the effect size. **(3)** *Substrate-recognition-site proximity* : minimum *Cα* distance from each variant’s mutated residue to any residue of the six substrate-recognition site (SRS) regions, with SRS boundaries taken from the universal selectivity-determining positions^33^ and mapped onto CYP2C9’s AlphaFold structure^13^ using SecStrAnnotator secondary-structure annotations,^34^ correlated with error by Spearman. **(4)** *Secondary structure*: structure elements assigned with SecStrAnnotator,^34^ with error compared across types by Kruskal–Wallis test.

#### 2.3.4. Clinical and pharmacogenetic benchmarking

We compared model-predicted activity against the held-out ClinVar and PharmVar labels (Section 2.1), which were excluded from training, using Spearman’s *ρ* between predicted activity and the ordinal function labels, where higher values denote greater retained function. PharmVar comparisons were additionally stratified by evidence level (definitive; definitive + moderate; all known-function alleles). As a baseline reference, we also correlated measured DMS activity directly against the same labels. Note the limited available data: only six CYP2C9 missense variants both carry a non-uncertain ClinVar label and are present in the DMS, and PharmVar evidence tiers range from *n* = 9 (definitive) to *n* = 27 (all known function) alleles.

#### 2.3.5. Substrate-resolved analysis

ClinPGx variant–drug annotations for all six CYPs were deduplicated to the best-evidence entry per (variant, drug) pair. A variant can exhibit substrate dependence only if it has been annotated against more than one drug, so the discordance rate was computed over the subset of variants carrying at least two drug annotations. A variant was called discordant if its function label was not identical across its drugs.

## 3. Results

### 3.1. AM fails systematically on CYPs

AM assigns CYP missense variants to its ambiguous pathogenicity class far more often than it does across the human proteome as a whole. Pooling all possible missense variants across the six CYPs (N=56,677 variants), 18.16% were classified as ambiguous, versus a proteome-wide baseline of 10.28% reported by AM^12^ — a 1.77-fold enrichment that is highly significant by a one-sided binomial test (*p <* 10^−16^) and consistent across all six CYPs examined (Fig. 1).

**Fig. 1.**
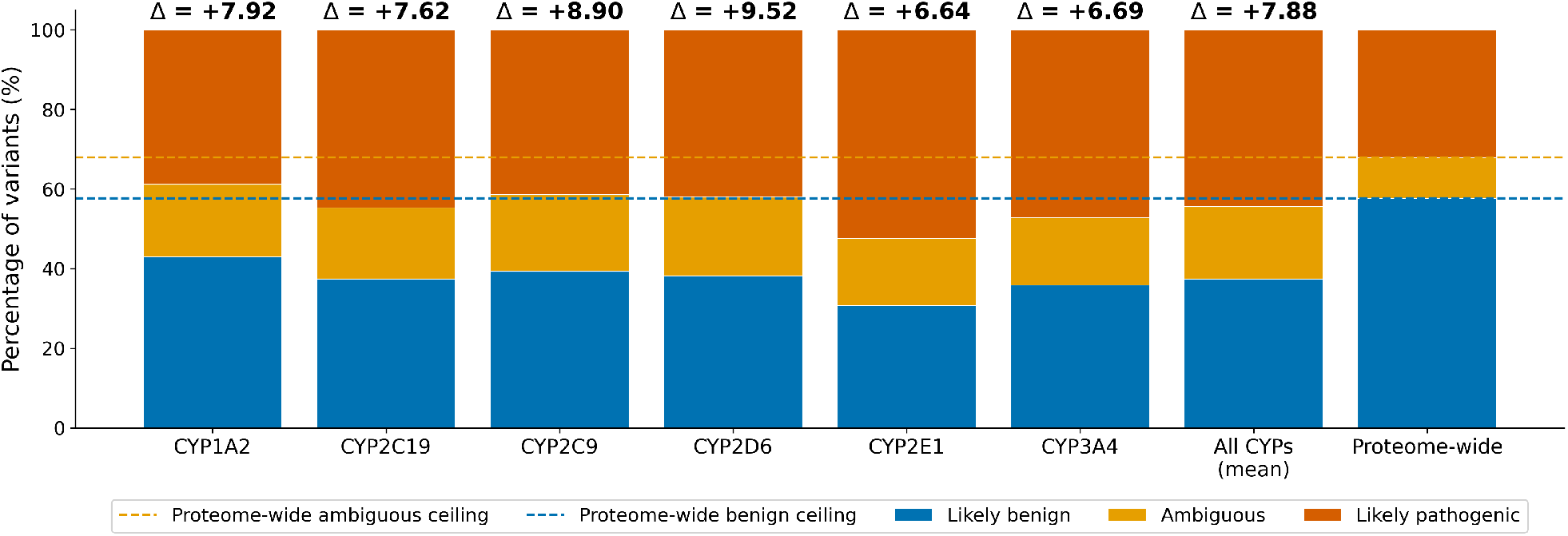
Rate of AM ambiguous pathogenicity labels in CYPs versus the proteome-wide baseline.^12^ Each Δ above a bar is the difference (in percentage points) between that CYP’s ambiguous fraction and the proteome-wide ambiguous fraction. Dashed lines mark the proteome-wide ambiguous and likely benign reference rates.

The correlation between AM’s pathogenicity score and measured activity is strong overall but uneven across classes: moderate within the likely benign (*ρ* = −0.354) and likely pathogenic (*ρ* = −0.423) classes and weak within the ambiguous class (*ρ* = −0.130) (Fig. S3). These correlations are negative because higher predicted pathogenicity tracks lower activity. To ask what AM is missing, particularly in the ambiguous class, we split the CYP2C9 DMS variants at the median abundance into destabilizing (low-abundance) and abundance-independent (normal/high-abundance) subsets, which reveals an abundance bias (Fig. 2). Across the full dataset, AM’s correlation with activity is strong for destabilizing variants (*ρ* = −0.568) but substantially weaker for abundance-independent variants (*ρ* = −0.357), a highly significant difference by an independent-samples Fisher *r*-to-*z* test (*z* = −8.99, *p <* 10^−18^). The two strata are matched on score range, IQR, and standard deviation, so the difference is not a range-restriction effect. Within the ambiguous class the gap is even more pronounced: AM retains signal for destabilizing ambiguous variants (*ρ* = −0.186, *p* = 1.5 *×* 10^−4^) but marginally at best for abundance-independent ones (*ρ* = −0.059, *p* = 0.234). The collapse is specific to this class; in the benign and pathogenic classes both abundance strata remain significant (*ρ* from −0.343 to −0.251, all *p <* 10^−14^). Consistent with this, AM’s score tracks cellular abundance directly, correlating with DMS abundance at *ρ* = −0.598 (Fig. S4), nearly as strongly as with activity itself (*ρ* = −0.643), despite AM being neither trained on abundance nor given it as input. AM therefore predicts the protein abundance component of variant effect well and the abundance-independent, catalytic component only weakly, pointing to a missing axis of activity independent of protein level.

**Fig. 2.**
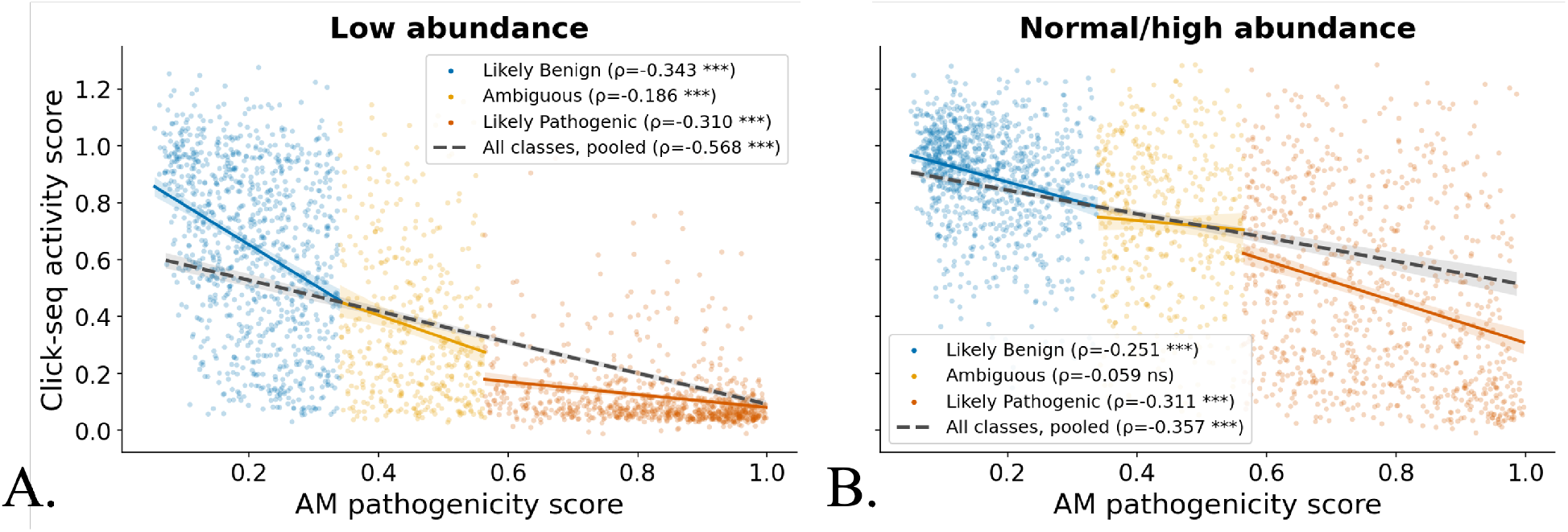
AM pathogenicity score versus CYP2C9 DMS click-seq activity, stratified by variant abundance (low abundance *N* = 2,210, **A**; normal/high-abundance *N* = 2,211, **B**). Each point is one missense variant, colored by AM class (likely benign, ambiguous, likely pathogenic); solid lines are ordinary least-squares fits within each class and the dashed line is the fit across the entire dataset, both with 95% confidence bands. Two-sided Spearman’s *ρ* is annotated per class and for the pooled data, where ^*∗∗∗*^*p <* 0.001; ns, not significant. Per-class variant counts are *n* = 952, 410, and 848 (likely benign, ambiguous, likely pathogenic) in (**A**) and *n* = 953, 410, and 848 in (**B**).

### 3.2. Biological priors only weakly explain AM prediction error

If AM’s failure had a structural or biochemical basis, biologically interpretable features should account for where its predictions go wrong. We tested four such features against AM’s absolute prediction error across the full dataset (*N* = 6,141). None explain the error well (Fig. S5).

Sequence position is unrelated to error (*ρ* = −0.002, *p* = 0.858; Fig. S5A). Substitution chemistry and secondary-structure type both reach significance on the full dataset (*H* = 19.954, *p* = 1.7 *×* 10^−4^; and *H* = 13.029, *p* = 0.023; Fig. S5B, D), albeit with very small effect sizes (*η*^2^ = 0.003 and 0.001, respectively). Each accounts for under 0.5% of the variance in prediction error, so significance here largely reflects statistical power at this sample size rather than a substantive effect.

Another structural feature with a defensible mechanistic rationale is proximity to the substrate-recognition sites, since CYP substrate selectivity is largely determined by the six SRS regions. SRS distance is the strongest of the four priors and is highly significant (*ρ* = −0.091, *p* = 8.6 *×* 10^−13^), with variants nearer the SRS regions carrying somewhat higher error (Fig. S5C). It nonetheless accounts for under 1% of the variance in error (*ρ*^2^ *≈* 0.008).

### 3.3. A CYP-specific ensemble is the most accurate activity predictor

Although AM and ESM-2 are strongly correlated overall (*ρ* = −0.797), their agreement is far weaker within the ambiguous class (*ρ* = −0.223) than within the benign (*ρ* = −0.488) or pathogenic (*ρ* = −0.615) classes (Fig. S3C). The two predictors thus diverge precisely where AM is least informative. Because AM and ESM-2 carry partly independent signal, most notably in the ambiguous class, we reasoned that combining the ESM-2 representation under protein-specific supervision with AM’s calibrated predictions could recover the catalytic axis AM misses. On a footing-matched comparison across an identical set of variants (*n* = 2,040, limited by a variant’s single-nucleotide reachability and dbNSFP coverage), a CYP2C9-specific KNN over ESM-2 embeddings, ensembled with calibrated AM and ESM-2 MMP, is the most accurate activity predictor we evaluate (overall *ρ* = 0.721; Table 1), exceeding AM (0.650) and ESM-2 (0.686) as well as every external baseline: the pharmacogene-tuned PharmGScore (0.604), the alignment-based SIFT (0.584), the supervised ensemble REVEL (0.539), and the function-trained DeMaSk (0.523). Because we did not test these differences for significance, the ensemble is nominally rather than demonstrably the best overall predictor.

**Table 1.** Predictive performance against CYP2C9 DMS activity: Spearman’s *ρ* between predicted and measured activity (out-of-fold, position-grouped 5-fold cross-validation), on a footing-matched set of *n* = 2,040 variants scored by all nine models. The protein-specific ensemble is the most accurate and exceeds every external baseline on equal footing.

| Model | Spearman $\rho$ | Paradigm |
| --- | --- | --- |
| AM + ESM-2 + KNN | **0.721** | protein-specific ensemble |
| AM + ESM-2 | 0.713 | ensemble |
| ESM-2 | 0.686 | protein language model (zero-shot) |
| AM | 0.650 | homology / pathogenicity |
| KNN (ESM-2 embeddings) | 0.625 | protein-specific |
| PharmGScore | 0.604 | pharmacogene-tuned |
| SIFT | 0.584 | alignment-based conservation |
| REVEL | 0.539 | supervised ensemble |
| DeMaSk | 0.523 | function-trained (DMS) |

The interpretable gain is concentrated where AM is weakest. Within the ambiguous class, AM alone reaches only *ρ* = 0.080, and adding it to ESM-2 changes nothing (0.341 for AM + ESM-2, identical to ESM-2 alone at 0.341) — AM contributes no usable ranking information there. The KNN alone reaches 0.353 and the full ensemble 0.399, a gain of +0.319 over AM alone; in comparison, the full ensemble improves on AM by +0.230 in the benign class and +0.098 in the pathogenic class, so the ambiguous class is where the ensemble changes most relative to AM’s own starting point (Table S1). These per-class correlations are computed on the full DMS set rather than on the footing-matched subset, since the external baselines are not defined for all variants; overall values therefore differ slightly from Table 1.

### 3.4. Predicted activity does not align with clinical or pharmacogenetic annotations

Recovering biochemical activity does not translate into reliable agreement with CYP2C9 clinical or pharmacogenetic function labels from ClinVar and PharmVar, respectively (Fig. 3A–B). Both label sets are ordinally encoded from no function to normal function, so agreement would appear as a positive correlation. Against ClinVar, no method reaches significance and the largest nominal correlations point the wrong way (*ρ* from −0.219 to −0.538, with corresponding *p* from 0.637 to 0.206; Fig. 3C). The same pattern holds against PharmVar under a high evidence bar: restricted to definitive-evidence alleles, every method is non-significant (*ρ* from −0.137 [SIFT] to −0.365 [KNN]), and admitting moderate evidence does not change this (Fig. 3D). Only when the threshold is relaxed to all alleles with any known-function label do several methods turn positive and significant (AM *ρ* = 0.505, *p* = 0.004; ESM-2 *ρ* = 0.399, *p* = 0.039; AM + ESM-2 *ρ* = 0.498, *p* = 0.008; AM + ESM-2 + KNN *ρ* = 0.450, *p* = 0.018;

**Fig. 3.**
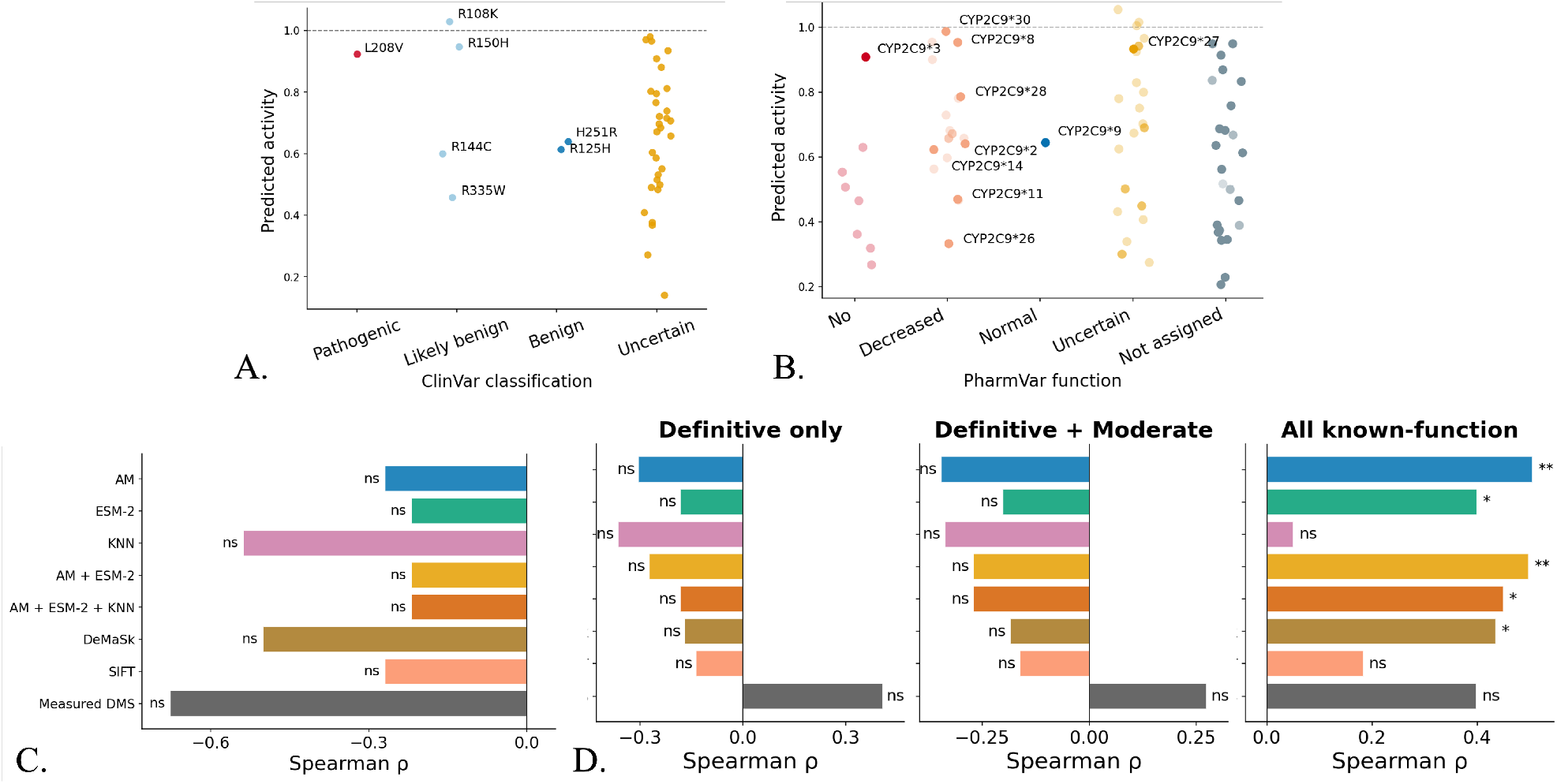
Predicted activity of CYP2C9 missense variants plotted against categorical labels from ClinVar and PharmVar. **A, B**. Predicted CYP2C9 activity (*y*-axis) versus ClinVar (**A**) or PharmVar (**B**) annotations (*x*-axis) for the AM + ESM-2 + KNN ensemble model. Each point is a single missense variant; select variants are labeled. In (**B**), marker opacity encodes PharmVar evidence level: fully opaque markers denote definitive-evidence alleles and progressively more transparent markers denote moderate- and limited-evidence alleles. **C, D**. Spearman correlation of predicted activity with ClinVar (**C**) or PharmVar (**D**) annotation, by model. Significance is indicated per bar, where ^*∗*^*p <* 0.05; ^*∗∗*^*p <* 0.01; ^*∗∗∗*^*p <* 0.001; ns, not significant. In (**D**), separate correlations are shown at the three PharmVar evidence thresholds (definitive only [*n* = 9], definitive + moderate [*n* = 11], all known function [*n* = 27]). The near-uniform absence of significance is the result: neither any method nor the measured assay separates from the rest.

DeMaSk *ρ* = 0.436, *p* = 0.016). That tier adds the limited-evidence alleles and therefore the noisiest labels, and the effect does not survive at either stricter threshold, so we read it as weak evidence rather than recovered agreement.

Even measured DMS click-seq activity reaches no significant agreement at any threshold (ClinVar *ρ* = −0.676, *p* = 0.140; PharmVar definitive *ρ* = 0.408, *p* = 0.363; all known function *ρ* = 0.399, *p* = 0.059). Every method we evaluate hits the same ceiling, and so does the experimentally measured assay itself. The limit is therefore not that some implementations read homology poorly while others perform better; the best use of homology we can construct sits at the ceiling with the rest. What thus fails is the shared target: a mismatch between what these methods predict and what the clinical labels encode.

These correlations rest on very few labeled variants, so individual annotations carry large leverage. CYP2C9*3 (I359L) is predicted to retain 71–91% of wild-type activity by every model, diverging not only from its “No Function” PharmVar label (assigned at the highest tier of evidence) but also from its measured DMS activity of 45%. L208V, ClinVar’s sole CYP2C9 variant carrying a “pathogenic” clinical-significance assertion, shows the opposite pattern: its predicted and measured activities agree (0.92 vs. 0.93) and both contradict the clinical label. DMS activity, clinical pathogenicity, and pharmacogenetic function are therefore related but distinct constructs, and accuracy against a biochemical assay does not guarantee concordance with clinical or pharmacogenetic annotations.

### 3.5. Variant effects depend on the substrate

In ClinPGx, the same variant is annotated with different functional consequences for different drug substrates, across all six CYPs examined (Fig. 4 shows CYP2C9 and CYP2D6; all six are in Fig. S6). Pooling across the six genes and restricting to variants with at least two drug annotations — the only variants for which substrate dependence is observable at all — 23 of 32 (71.9%) carry a function label that differs by drug. Restricting further to annotation pairs sharing an endpoint type leaves discordance essentially unchanged, at 20 of 28 (71.4%). Although sparse per-variant annotations leave formal testing underpowered, a discordance rate this high indicates that substrate identity, fixed by construction in the DMS assay and absent altogether from every model evaluated above, carries variant-effect information that a single per-variant activity value cannot represent.

**Fig. 4.**
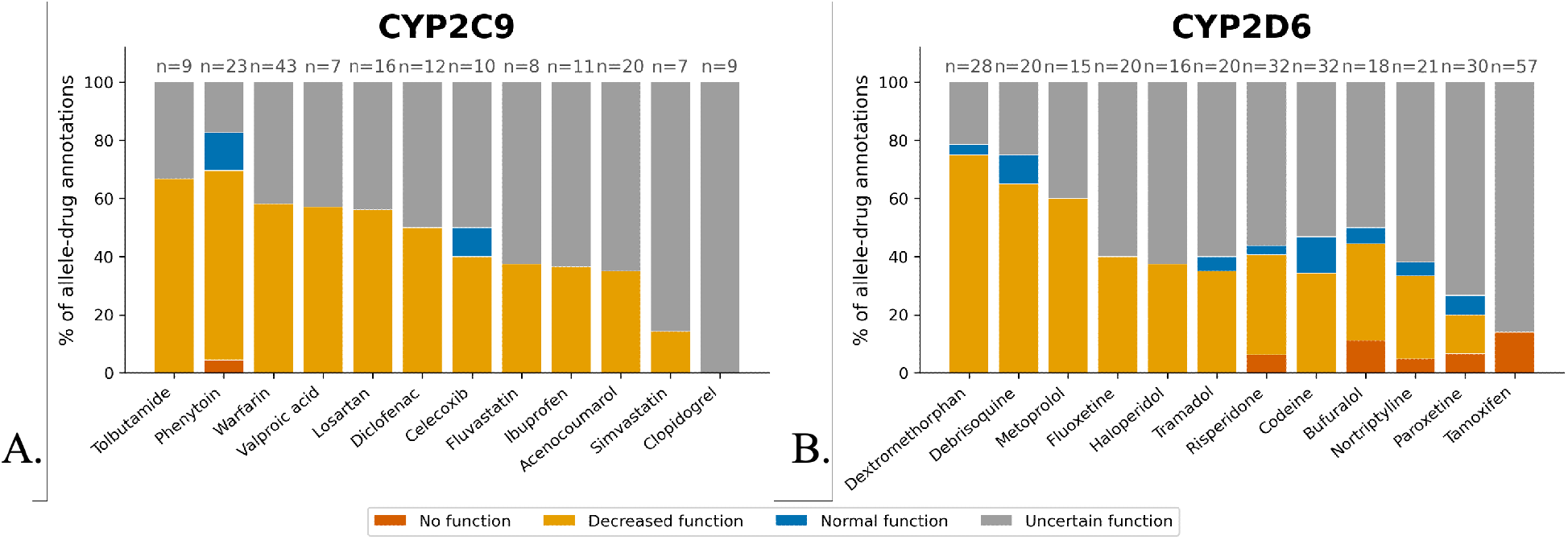
Substrate-resolved variant effects. Stacked bar charts showing, for two representative CYP genes (CYP2C9 (**A**) and CYP2D6 (**B**)), the distribution of ClinPGx function annotations (no function, decreased function, normal function, and uncertain function) across the top 12 drug substrates for each CYP (*x*-axis). Each bar is one drug and shows the percentage of allele–drug annotations assigned each function label (*y*-axis, % of allele–drug annotations, 0–100%), where one annotation is a single allele–drug pair. The number of annotations (*n*) is given above each bar. The complete version for all six CYPs examined is Fig. S6.

## 4. Discussion

Using CYP pharmacogenes, we found that homology-based prediction fails systematically. AM reads the abundance axis of variant effect and misses an abundance-independent, catalytic one, most completely within the ambiguous class that CYPs are enriched for — a residual that sequence position, substitution chemistry, substrate-recognition-site proximity, and secondary structure do not explain. A protein-specific ensemble drawing on homology-derived features (KNN on ESM-2 embeddings, AM, and ESM-2 MMP) is nominally the most accurate activity predictor we evaluate, exceeding every generic, function-trained, and pharmacogene-tuned baseline. Yet it too fails against clinical and pharmacogenetic labels, as do all of those baselines and the measured assay itself. The breakdown we report is therefore not that these models read homology badly, which better use of the same representation would repair; it is that the best available use of homology-derived information reaches a ceiling short of what the clinical application of pharmacogenomics requires. We further show that the same variant is annotated with different consequences for different drug substrates across CYPs in ClinPGx, pointing to substrate identity as a determinant of CYP activity that no current predictor captures.

### 4.1. Why homology breaks down on pharmacogenes

Homology-based predictors succeed when evolutionary tolerance is a good proxy for functional consequence. For fold stability it usually is: a substitution that destabilizes the protein tends also to be evolutionarily disfavored, so scores derived from the residue distribution across homologs track it. Catalytic activity toward a specific substrate is different — it can be tuned by active-site substitutions that leave the fold intact and that selection, acting on xenobiotic breadth rather than on a single conserved catalytic output, need not have constrained.^3,4,6^ Our abundance decomposition makes this concrete: AM’s signal survives among destabilizing variants and vanishes among abundance-independent ones, exactly the split between the abundance axis it can see and the catalytic axis it cannot. This agrees with reports in other proteins that AM scores overlap with computed destabilization energies (ΔΔ*G*) most strongly at buried residues,^35^ and with broader evidence that generic pathogenicity scores carry structural rather than catalytic information.^36,37^

### 4.2. Function is multi-dimensional, and existing ground truth reflects this

A predictor is only as well posed as its target, and our results make explicit that the resources used to define variant effect are not interchangeable: biochemical activity toward a probe substrate (DMS), disease causation (ClinVar), pharmacogenetic function (PharmVar), and substrate-resolved consequence (ClinPGx) answer different questions. That a model can reproduce measured activity closely yet not align with clinical labels is therefore expected rather than anomalous once function is recognized as multi-dimensional. The recent literature independently supports this. A DMS of CYP2C19 uncovered a substrate-specificity–abundance tradeoff within a single assay,^26^ showing that even one experiment’s notion of function is substrate-contingent; the difficulty of assigning substrate-dependent allelic effects is a standing clinical challenge;^38^ and CYP2C9 activity depends even on cofactor context.^39^ The ceiling that variant-effect predictors (VEPs) encounter is therefore as much definitional as methodological, since the target itself is underspecified when a CYP variant’s effect is collapsed to a single substrate-agnostic number.

### 4.3. Relationship to prior computational approaches

Our protein-specific recovery of activity signal fits an active trend of adapting general predictors to narrower problems, from fine-tuning protein language models on DMS data^40–42^ to pharmacogene-specific models that outperform off-the-shelf tools.^19,22–25^ Using established predictors, including the DMS-derived DeMaSk^18^ and the pharmacogene-calibrated PharmGScore,^19^ alongside our CYP2C9-specific ensemble, we show that adaptation to functional data improves activity accuracy without moving a model toward the clinical target, pointing to a ceiling that adaptation alone cannot cross.

While some models do incorporate substrate identity, they remain narrow proofs of principle. Site-of-metabolism and enzyme-kinetics models predict how the wild-type enzyme handles a given drug,^43–48^ but do not predict how a genetic variant alters metabolism of a specific substrate. Conversely, the widely used VEPs are substrate-agnostic. Work that does combine the two remains protein- or task-specific.^49,50^ To our knowledge, no widely used, generalizable VEP conditions on the substrate.

### 4.4. Toward substrate-conditioned variant-effect prediction

We therefore propose substrate identity as the leading candidate for the non-homologous variable that current predictors omit, and substrate-conditioned prediction as the path beyond homology. Rather than mapping a variant to a scalar, such a model would predict function for a (variant, substrate) pair, most naturally as substrate-specific kinetic parameters of the kind that enzyme-kinetics models already target for wild-type enzymes.^47,48^ Direct biochemistry is consistent with this hypothesis: a single active-site substitution can shift CYP2C9 metabolism in opposite directions across substrates. F476W raises (S)-warfarin catalytic efficiency five-fold while lowering (S)-flurbiprofen twenty-fold,^51^ and active-site mutations reorder turnover of warfarin, flurbiprofen, and phenytoin differently.^52^ These findings corroborate the substrate discordance we observe statistically in ClinPGx: for a multi-substrate enzyme, a variant’s effect is a function of the substrate, which is exactly what a substrate-conditioned predictor must learn. With the biological rationale thus established, the principal remaining obstacle is data: substrate-resolved measurements are sparse, so we present this as a direction rather than a result, though multi-substrate proof-of-concept studies suggest it is increasingly tractable.^53^ We consider two additional hypotheses as well. The first is that the missing axis is not substrate identity but cellular and physiological context. Expression level, redox-partner and cofactor availability, membrane environment, and compensating activity from other CYPs all modulate the phenotype a variant produces in vivo, and none is captured by a sequence-based score or by a single-cell-line assay. CYP2C9 activity is known to depend on cofactor context,^39^ and this hypothesis could likewise explain the failure against clinical labels. The second is that the gap is substantially label-side. PharmVar and ClinVar labels are assigned from heterogeneous evidence of varying quality, are frequently driven by a single early study, and encode clinical judgment rather than a biochemical quantity, so part of the ceiling may reflect noise in the annotations rather than a missing biological variable. We regard substrate dependence as the most likely hypothesis, for three reasons: it is directly observed in the annotation data, it is corroborated by direct CYP2C9 biochemistry,^51,52^ and it predicts precisely the abundance-independent, catalysis-specific blind spot our decomposition finds.

The clinical motivation also aligns with this framing. Genotype-guided prescribing is inherently drug-specific,^9,10^ so a substrate-conditioned predictor targets exactly what the clinic requires, whereas a single activity score does not. The most promising route beyond homology here thus runs less through better similarity search than through enriching the definition of function, so that prediction is conditioned on substrate in addition to sequence conservation.

### 4.5. Limitations

Several limitations bound these conclusions. Our class-stratified analyses inherit AM’s own class definitions: the ambiguous class is a band of AM’s score axis fixed by the thresholds AM applies proteome-wide, not an independently motivated partition of CYP2C9 variants, so results reported are conditional on where those thresholds fall. Our activity and abundance benchmarks rest on a single DMS study of one gene;^8^ whether the abundance bias generalizes to other CYPs awaits comparable paired data. The recovered accuracy comes from a protein-specific, recalibrated model and does not generalize beyond CYP2C9. Combining biochemical, clinical, and pharmacogenetic data is confounded by differences in measurement, thresholds, expression system, and evidence level. The clinical and pharmacogenetic comparisons rest on few labeled variants, so correlations are largely non-significant, sensitive to individual labels, and unstable in sign across evidence tiers. Finally, the sparse substrate-resolved annotations in ClinPGx establish that substrate-dependent effects occur for a meaningful fraction of variants but leave their overall prevalence underpowered to estimate.

## Acknowledgements

This work is supported by NIH GM153195, NIH NLM F31LM014646, Burroughs Wellcome Fund IRSA 1074128, and a Stanford Medical Scholars award. We thank Stanford Research Computing for providing computational resources that contributed to this research.

