## Supplementary Materials for "Homology-Based Variant-Effect Predictors Break Down on Cytochrome P450 Pharmacogenes^*^"

Supplementary materials accompanying the main text.

© 2026 The Authors. Open Access chapter published by World Scientific Publishing Company and distributed under the terms of the Creative Commons Attribution Non-Commercial (CC BY-NC) 4.0 License.

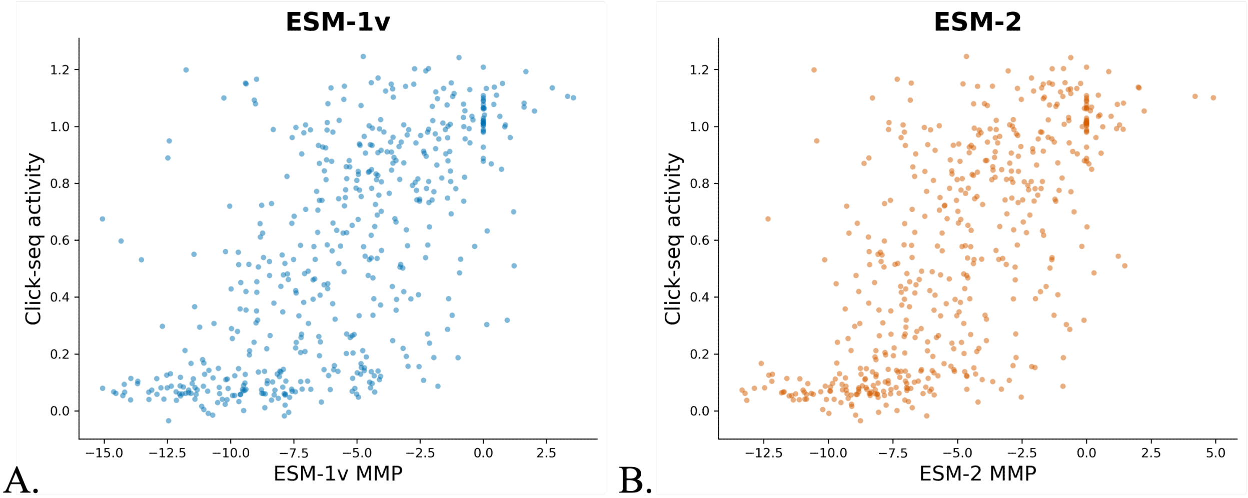


**Fig. S1**. ESM-1v versus ESM-2 masked marginal probability (MMP) as a predictor of CYP2C9 activity. ESM-1v (**A**) and ESM-2 (**B**) MMP against normalized CYP2C9 DMS click-seq activity for a shared set of 500 randomly sampled variants. Two-sided Spearman's rank correlation: ESM-1v $\rho = 0.657$ ($p = 3.7 \times 10^{-63}$); ESM-2 $\rho = 0.697$ ($p = 5.3 \times 10^{-74}$). ESM-2 correlates with DMS activity more strongly than ESM-1v, a significant difference by a Steiger–Meng test for dependent correlations ($z = -2.92$, two-sided $p = 0.0035$). On this basis, ESM-2 was adopted as the masked marginal predictor throughout.

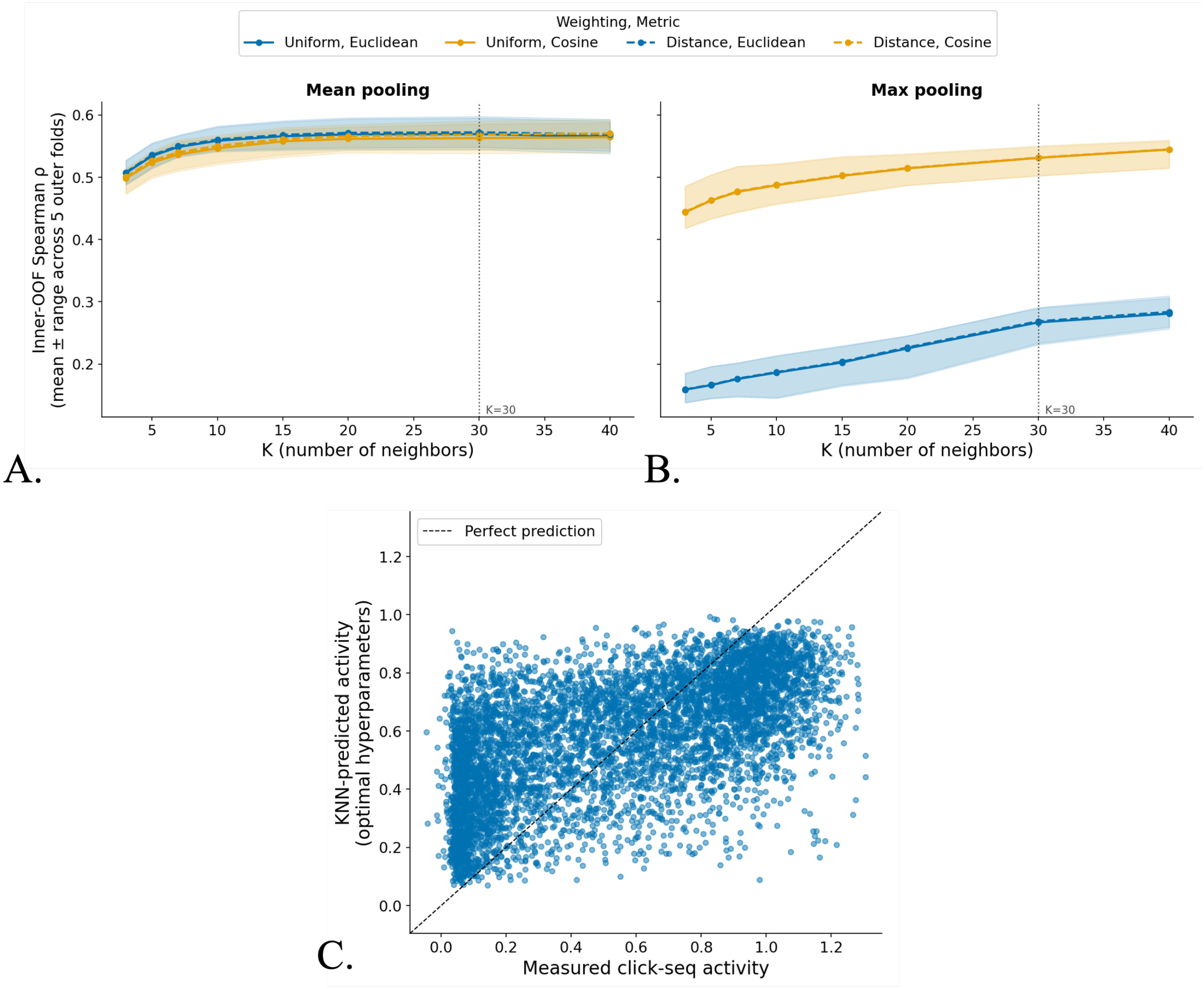


**Fig. S2**. KNN hyperparameter selection. Hyperparameters of the ESM-2 embedding KNN regressor predicting normalized CYP2C9 click-seq activity were selected by a unified grid search over pooling of the final-layer per-residue embeddings (mean, max) $\times$ neighbor weighting (uniform, distance-weighted) $\times$ distance metric (Euclidean, cosine) $\times$ number of neighbors $K \in \{3, 5, 7, 10, 15, 20, 30, 40\}$. Selection is nested inside evaluation: an outer 5-fold position-grouped split holds out each fold's residue positions entirely, and within each outer training partition a 3-fold position-grouped inner split scores all configurations. The inner winner is refit on the full outer training partition and applied to the held-out outer fold. **A, B.** Inner out-of-fold Spearman $\rho$ against $K$ for each weighting $\times$ metric combination, under mean (**A**) and max (**B**) pooling; lines are the mean across the five outer folds and shaded bands the range. The dotted vertical line marks $K = 30$. Mean pooling and distance weighting were selected in all five outer folds, while the selected $K$ and distance metric varied within a narrow region of the grid ($K \in \{30, 40\}$; Euclidean in four of five folds). The configuration selected most often across outer folds (mean pooling, distance-weighted averaging, Euclidean metric, $K = 30$) was used in all subsequent analyses. **C.** Out-of-fold predicted activity versus measured click-seq activity for this fixed hyperparameter configuration with the dashed line indicating perfect prediction ($y = x$); Spearman $\rho = 0.597$.

A.
B.
C.

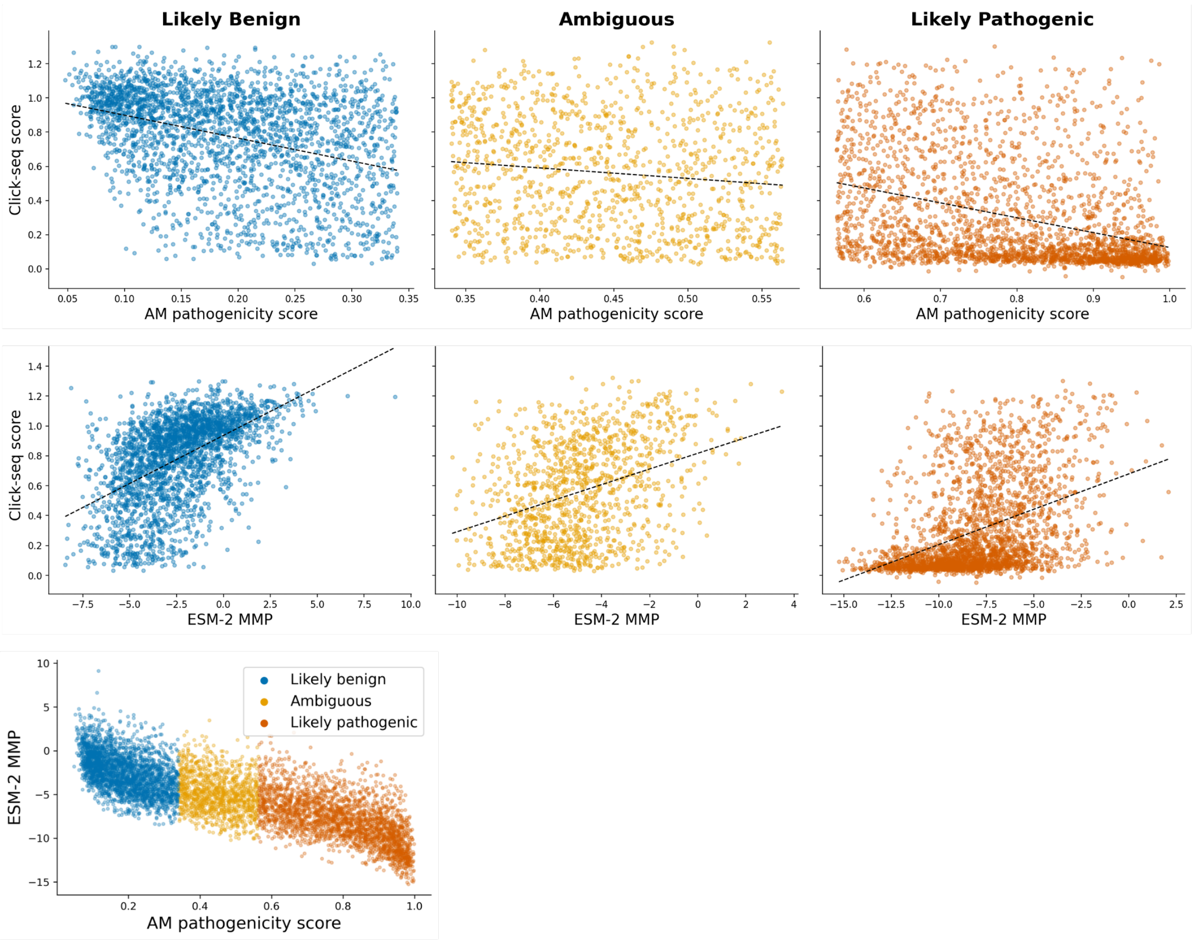


**Fig. S3**. AM pathogenicity score (**A**) and ESM-2 MMP (**B**) versus CYP2C9 DMS click-seq activity, stratified by AM class (likely benign, ambiguous, and likely pathogenic). Each point is one missense variant, and dashed lines indicate ordinary least-squares linear fits. For AM (**A**), the score–activity relationship is moderate within the likely benign ($\rho = -0.354$) and likely pathogenic ($\rho = -0.423$) classes and weak within the ambiguous class ($\rho = -0.130$). ESM-2 shows the same pattern: likely benign $\rho = 0.544$, ambiguous $\rho = 0.343$, and likely pathogenic $\rho = 0.446$ (**B**). **C.** ESM-2 MMP versus AM pathogenicity score, with points colored by AM class; pooled $\rho = -0.797$, likely benign $\rho = -0.488$, ambiguous $\rho = -0.223$, likely pathogenic $\rho = -0.615$.

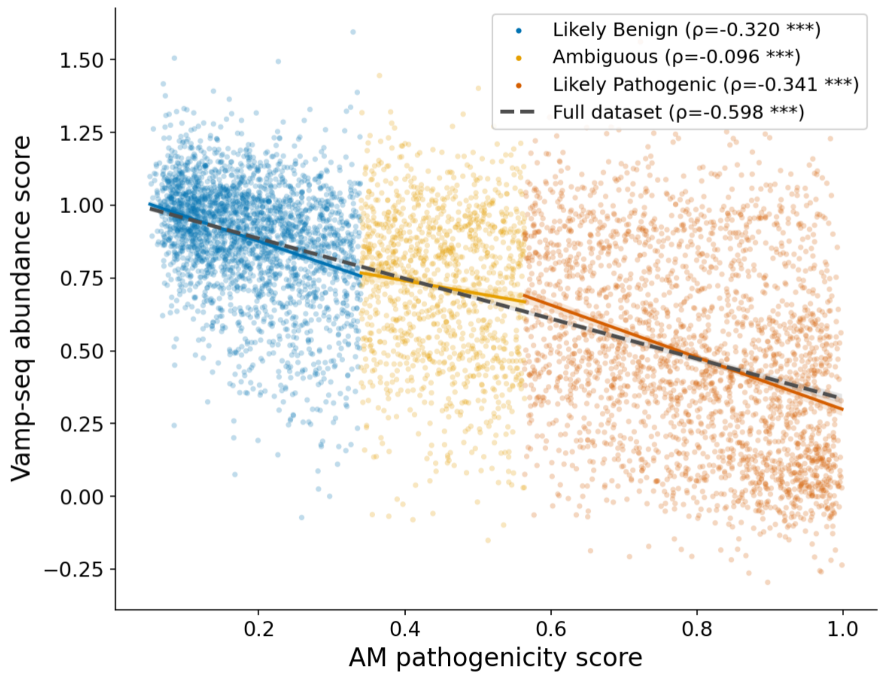


**Fig. S4**. AM pathogenicity score versus CYP2C9 DMS vamp-seq abundance; points are colored by AM class, solid lines are within-class least-squares fits, and the dashed line is the pooled fit. AM tracks abundance ($\rho = -0.598$, $p < 10^{-300}$) about as closely as it tracks catalytic activity ($\rho = -0.643$), even though abundance is a biophysical property AM was neither trained on nor given as input. The negative sign reflects higher predicted pathogenicity tracking lower abundance. As with activity, the relationship is moderate within the benign ($\rho = -0.320$) and pathogenic ($\rho = -0.341$) classes and weak within the ambiguous class ($\rho = -0.096$). $^{***}p < 0.001$.

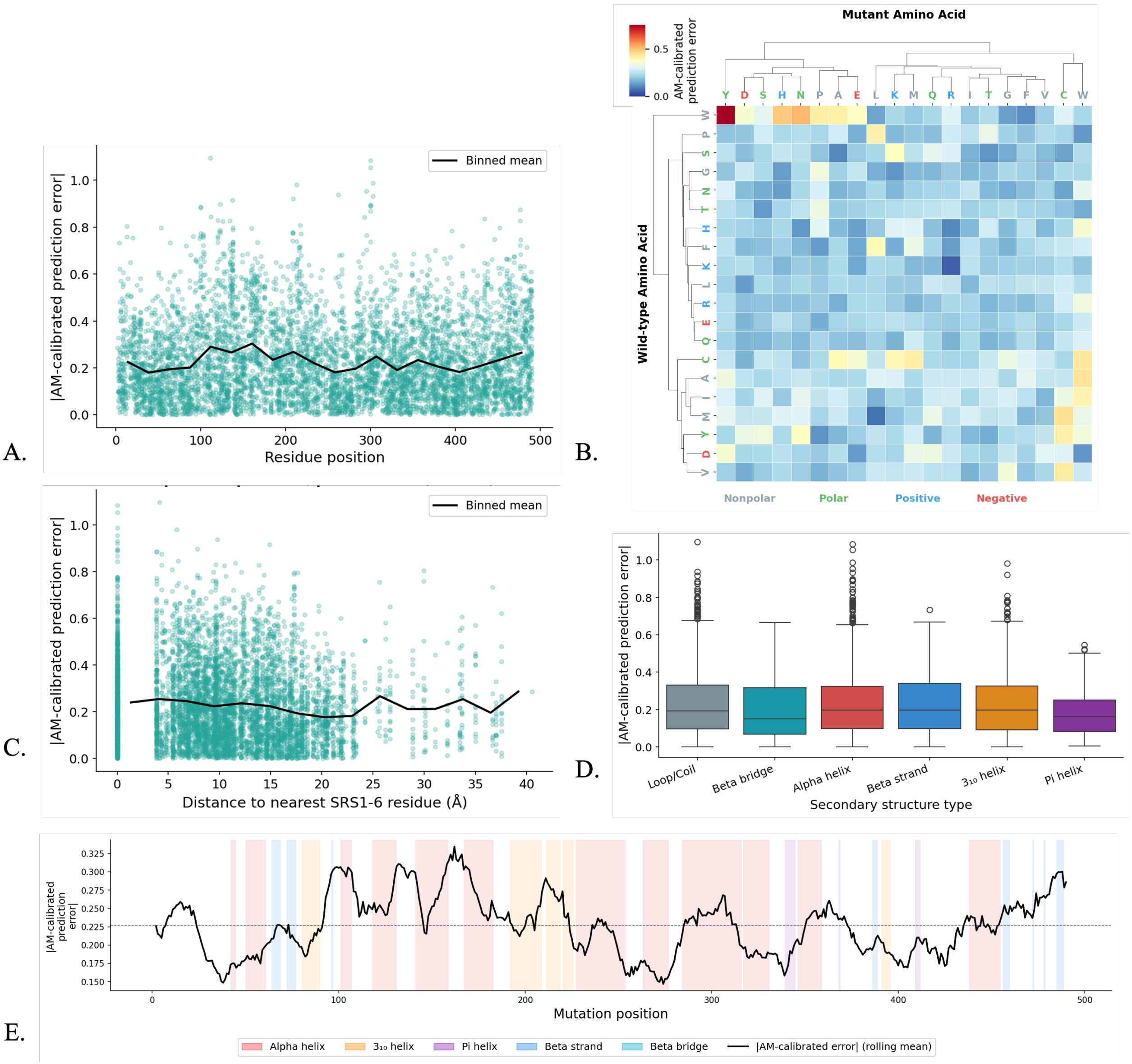


**Fig. S5**. Structural and biochemical priors versus AM prediction error for CYP2C9 missense variants ($N = 6{,}141$). Prediction error is the absolute residual between AM's calibrated activity prediction and measured click-seq activity. **A.** Error versus residue position; each point is one missense variant and the solid line is the binned mean (two-sided Spearman $\rho = -0.002$, $p = 0.858$). **B.** Mean error per wild-type → mutant amino-acid substitution, with rows and columns ordered by hierarchical clustering (Ward linkage) and labels colored by biochemical group (nonpolar, grey; polar, green; positive, blue; negative, red); error differs across wild-type residue groups by Kruskal–Wallis test ($H = 19.954$, $p = 1.7 \times 10^{-4}$, $\eta^2 = 0.003$). **C.** Error versus minimum C$\alpha$ distance to any residue of the six substrate-recognition site regions, with the binned mean overlaid ($\rho = -0.091$, $p = 8.6 \times 10^{-13}$). **D.** Error by secondary-structure type assigned with SecStrAnnotator; boxes show the median and interquartile range, whiskers extend to 1.5×IQR, and points beyond are plotted individually ($H = 13.029$, $p = 0.023$, $\eta^2 = 0.001$). **E.** Error along the CYP2C9 sequence (15-residue rolling mean), with secondary-structure elements shaded by type; the dashed line marks the protein-wide mean error.

**Table S1**. Predictive performance of individual models and ensembles against CYP2C9 DMS activity, on the full click-seq set ($N$ = 6,141). Spearman's $\rho$ between predicted and measured activity, reported overall and within each AM class, for AM, ESM-2, KNN, and the AM + ESM-2 and AM + ESM-2 + KNN ensembles (out-of-fold, position-grouped 5-fold cross-validation).

| Model | Overall $\rho$ | Benign $\rho$ | Ambiguous $\rho$ | Pathogenic $\rho$ |
|---|---|---|---|---|
| AM | 0.637 | 0.338 | 0.080 | 0.400 |
| ESM-2 | 0.677 | 0.543 | 0.341 | 0.441 |
| KNN | 0.597 | 0.440 | 0.353 | 0.342 |
| AM + ESM-2 | 0.698 | 0.551 | 0.341 | 0.475 |
| AM + ESM-2 + KNN | **0.713** | **0.568** | **0.399** | **0.498** |

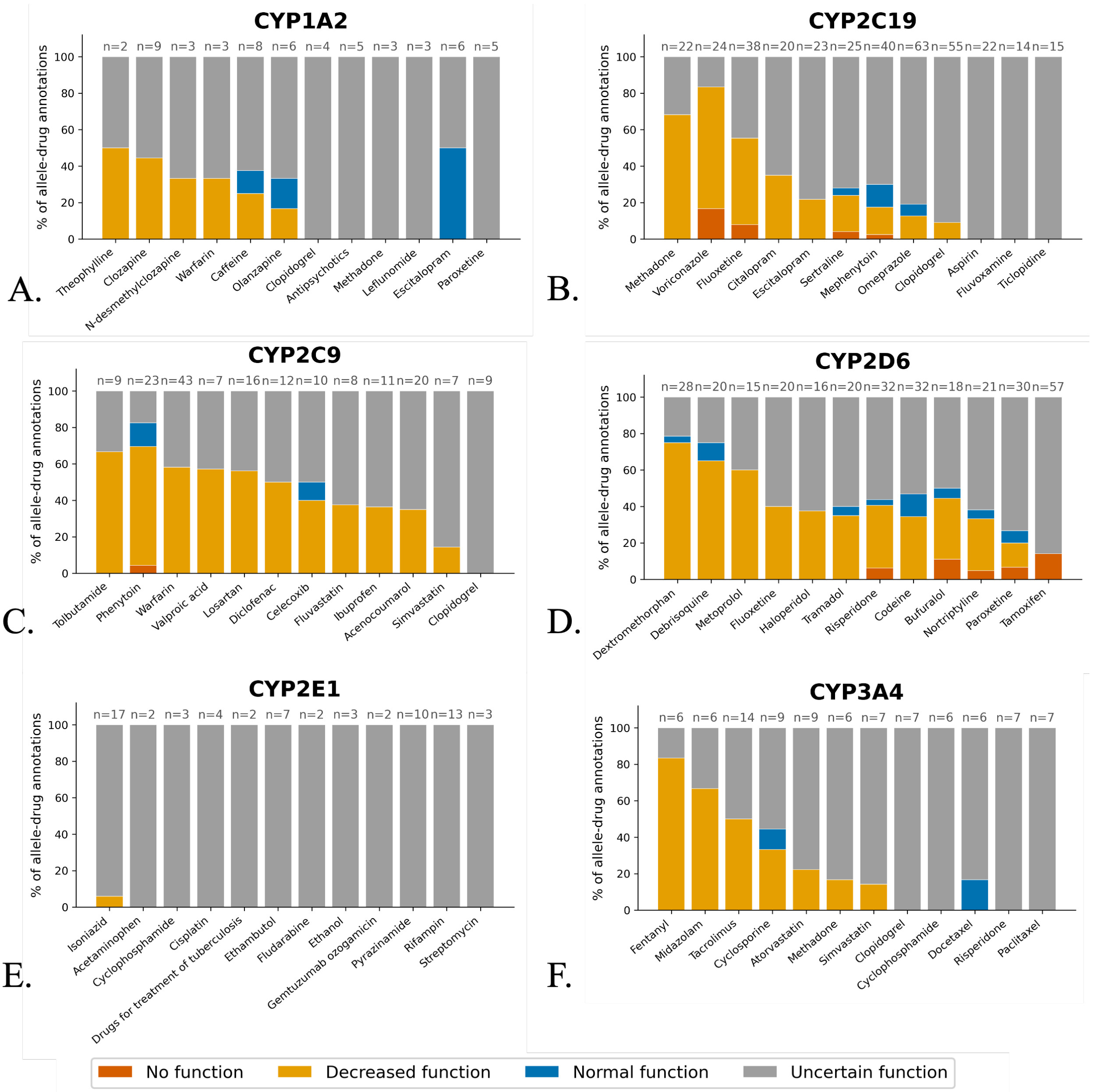


**Fig. S6**. Substrate-resolved variant effects across six CYPs from ClinPGx. Stacked bar charts showing, for each of the six CYP genes (CYP1A2, CYP2C19, CYP2C9, CYP2D6, CYP2E1, CYP3A4; **A–F**), the distribution of ClinPGx function labels (no function, decreased function, normal function, and uncertain function) across the top 12 drug substrates for each CYP ($x$-axis). Each bar is one drug and shows the percentage of allele–drug annotations assigned each function label ($y$-axis, % of allele–drug annotations, 0–100%), where one annotation is a single allele–drug pair. The number of annotations ($n$) is given above each bar. This is the six-gene extension of Fig. 4 of the main text.
